# Phospholipid tail asymmetry allows cellular adaptation to anoxic environments

**DOI:** 10.1101/2022.08.04.502790

**Authors:** Luca Panconi, Chris D Lorenz, Robin C May, Dylan M Owen, Maria Makarova

## Abstract

Membrane biophysical properties are critical to cell fitness and depend on unsaturated phospholipid acyl tails. These can only be produced in aerobic environments since eukaryotic desaturases require molecular oxygen. This raises the question of how cells maintain bilayer properties in anoxic environments. Here, we demonstrate the existence of an alternative pathway to regulate membrane fluidity that exploits phospholipid acyl-tail length asymmetry, replacing unsaturated species in the membrane lipidome. We show that the fission yeast, *S. japonicus*, which can grow in aerobic and anaerobic conditions, is capable of utilizing this strategy whereas its sister species, the well-known model organism *S. pombe*, cannot. The incorporation of asymmetric-tailed phospholipids might be a general adaptation to hypoxic environmental niches.

**One-Sentence Summary:** In anoxic environments, saturated asymmetric acyl-tailed phospholipids can replace unsaturated ones to maintain membrane physical properties.

## Main Text

The biophysical properties of cell membranes are critical for cell function and survival in varying environmental conditions. Cells use their metabolic capacity to generate lipidomes that optimise membrane properties and thereby adapt to changing environmental niches. This is because it is membrane lipids that predominantly define membrane properties such as fluidity, viscosity, bending rigidity and lateral diffusion (*1*). These properties in turn affect macroscale membrane activities such as curvature, fusion, fission and the lateral distribution of membrane proteins (*1*, *2*) Membrane properties are strongly defined by the hydrophobic core of the lipid bilayer, particularly by the phospholipid acyl tail composition. Fully saturated acyl tails form a tightly packed membrane with low lateral diffusion and, in combination with sterols, form the so-called liquid-ordered phase. Alternatively, when acyl tails contain a double bond, a kink is introduced into the structure which prevents tight lipid packing and forms a liquid-disordered phase with high lateral diffusion (*3*–*5*).

Desaturases are enzymes which introduce double bonds into acyl moieties in a process that is dependent on oxygen. Their di-iron centre is activated by molecular oxygen to remove two hydrogens from the acyl chain and generate a double bond (*6*, *7*), making oxygen a factor regulating membrane composition and properties. While some prokaryotic organisms can generate double bonds in the absence of oxygen during the synthesis of an acyl tail, this ability is not found in eukaryotes (*8*). The question therefore arises of how eukaryotic cells maintain their membrane properties in anaerobic conditions when they are unable to produce unsaturated-tail membrane phospholipids.

The fission yeast *Schizosaccharomyces japonicus* is known to grow in both aerobic and anaerobic conditions and, compared to its well-studied relative *Schizosaccharomyces pombe*, has a demonstrated plasticity for generating diverse phospholipid acyl tail profiles. Recently it has been shown that the *S. japonicus* lipidome contains high quantities of asymmetric acyl chain phospholipids with 18:0 and 10:0 carbon atoms as the most predominant configuration (*9*). This asymmetry in acyl tails is not exclusive to *S. japonicus*, having also been identified in *S. cerevisiae*, suggesting that these structural phospholipids could be more abundant than previously recognised (*10*). Our earlier work using artificial membrane systems has shown that these asymmetric phospholipids are able to replace unsaturated lipids to maintain the biophysical properties of simple, artificial membrane systems (*11*). We therefore hypothesise that the ability of *S. japonicus* to produce these asymmetric phospholipids may be behind its adaptation to anaerobic environments.

Here, we show that in the absence of oxygen, these asymmetric acyl tails fully replace unsaturated moieties and allow cells to maintain membrane properties to insure cell fitness. Deletion of the membrane bound desaturase ole1 Δ9 resulted in a similar shift toward fully saturated, asymmetric lipids. At 37 °C, neither the lack of oxygen not the desaturase deletion affected cell fitness, suggesting that saturated, asymmetric lipids can substitute for their unsaturated counterparts. However, this compensation was only possible at physiologically high temperatures, since cells grown at 24 °C exhibited growth defects. Molecular dynamics simulations and advanced microscopy using environmentally-sensitive probes traced this effect to an inability of asymmetric lipids to maintain membrane fluidity at these lower temperatures. Therefore, while asymmetric lipids may have evolved as a strategy for organisms to maintain membrane fluidity in the absence of the unsaturated lipids that molecular oxygen allows, this strategy is only partially successful and fails at low temperatures.

## Results

### Lack of oxygen induces lipidome rewiring in S. japonicus

To investigate how the membrane lipidome adapts to anaerobic growth in *S. japonicus*, we first assessed the growth pattern of *S. japonicus* compared to *S. pombe*. To fully deplete lipids that could have been synthesised in the presence of oxygen we grew cells in an anaerobic cabinet for 48 h in a synthetic mineral medium and reinoculated cells into fresh oxygen-free medium every 24 h (Fig. 1 A). After 48 h in anaerobic conditions, we measured cell growth and confirmed that *S. japonicus* growth rates were similar in both aerobic and anaerobic conditions (Fig. 1 B, C). In contrast, *S. pombe* showed complete growth suppression in anaerobic conditions (Fig. 1 C).

**Fig. 1.**
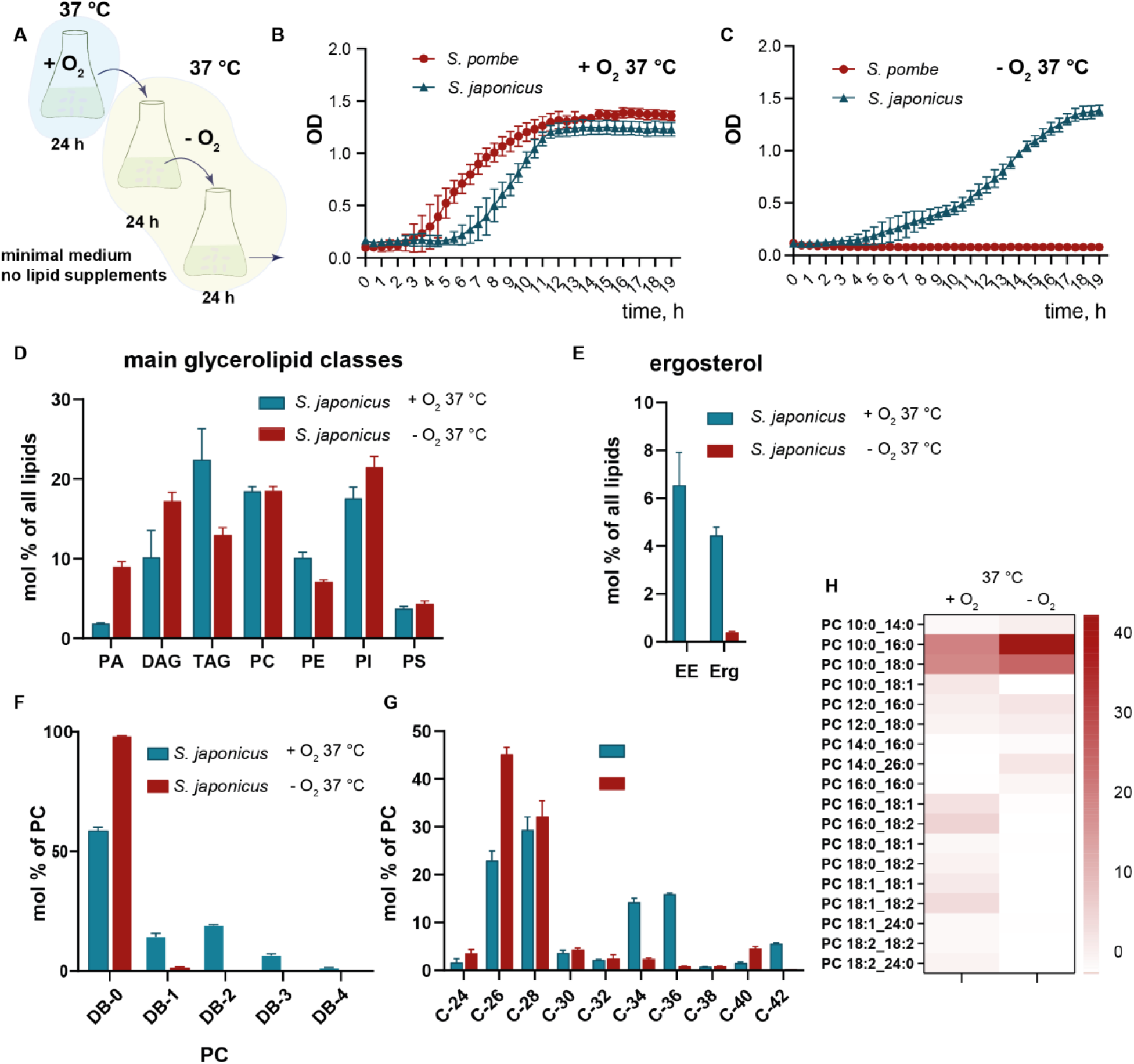
*S. japonicus* grown in anoxic conditions alter their membrane lipidome. (**A**) Scheme of growth experiment in the absence of oxygen, cells were grown in minimal medium conditions for 24 hours in the presence of oxygen to mid-log growth phase then transferred to the anaerobic chamber and diluted with minimal medium without oxygen. Cells were allowed to grow to mid-log phase in an anaerobic chamber before growth and lipidomic experiments. (B) Growth curves of *S. japonicus* and *S. pombe* in minimal medium with oxygen measured as a change of optical density (OD) at 600 nm. (C) Growth curves of *S. japonicus* and *S. pombe* in minimal medium without oxygen measured as a change of OD. (D) Relative abundance of main phospholipid and glycerolipid classes presented as a molecular percentage of all identified lipids in *S. japonicus* grown in the presence and absence of oxygen at 37 °C. (E) Relative abundance of ergosterol and ergosterol ester in *S. japonicus* grown in the presence and absence of oxygen at 37 °C. (F) Relative abundance of double bonds in acyl tails within the PC class in *S. japonicus* grown in normoxic and anoxic conditions. (G) Relative abundance of various carbon lengths in acyl tails in *S. japonicus* grown in normoxic and anoxic conditions. (H) A heat map showing molecular species composition within PC class in *S. japonicus* grown in normoxic and anoxic conditions. The colour bar represent molecular percentage of species within PC class.

To assess how *S. japonicus* adapts its lipid composition in the oxygen-free environment, we quantitatively analysed around 600 unique lipid species via shotgun mass spectrometry in cells grown for at least 48 h anaerobically in minimal medium at 37 °C, in comparison to cells that had been grown aerobically. The main significant differences in glycerolipid classes were observed in increased phosphatidic acid, diacylglycerol and decreased triglycerides during anaerobic growth. The main glycerophospholipid classes such as PC (phosphatidylcholine), PE (phosphatidylethanolamine), PI (phosphatidylinositol) and PS (phosphatidylserine) were only mildly altered in anaerobic conditions (Fig. 1 D). This shows that phospholipid head groups are not substantially affected by anaerobic growth conditions. We also identified differences in minor lipid classes - ceramides and lysophospholipids that were significantly increased when cells were grown in the absence of oxygen (table S1). As expected, in the prolonged absence of oxygen *S. japonicus* also depleted ergosterol together with esterified ergosterol (Fig. 1 E).

Despite these minor changes, the main adaptation to anaerobic growth in *S. japonicus* was in the lipid hydrophobic chain make-up - length and saturation (Fig. 1 F-H). The lack of oxygen-induced major rewiring in acyl chain composition in all identified classes of phospholipid, with unsaturated tails replaced by fully saturated counterparts (Fig. 1F - for PC, the distribution of lipid species in other classes is shown in fig. S1 and table S2). This was accompanied by a decrease in the average acyl tail’s length with lipid species that had a total of 34 or 36 carbon atoms in both tails being replaced by lipids mainly containing 26 carbon atoms (Fig. 1G). Previously, it is been shown that the *S. japonicus* lipidome is naturally enriched in asymmetric acyl tail lipids with a total carbon length of 26 or 28 atoms and with tails 18:0/16:0 and 10:0 as the most predominant configuration (*9*). We confirmed that these 26-carbon tail lipid therefore represent the asymmetric 16:0,10:0 species (Fig. 1H- for PC, the distribution of lipid species in other classes is shown in fig. S1). Lysophospholipids, ceramides, diacylglycerol and triacylglycerols showed similar changes (table S2).

Overall then, *S. japonicus* is able to grow normally in anaerobic conditions whereas *S. pombe* is not. The lack of oxygen induces dramatic changes in the *S. japonicus* lipidome with a complete loss of acyl-tail unsaturation and a dramatic increase in asymmetric phospholipids, primarily with saturated tails containing 16 and 10 carbons, respectively.

### S. japonicus upregulates asymmetric lipids specifically due to the absence of unsaturated acyl tails

The main desaturase in fission yeast that introduces the double bond into acyl moieties is the Ole1 Δ9 desaturase which has been shown to require molecular oxygen (*12*). We hypothesised that the increase in asymmetric lipids is a mechanism for the cell to compensate for the inability to synthesise unsaturated acyl tails in anaerobic conditions. To test this, we generated a mutant *S. japonicus* strain that lacks *ole1 Δ9* desaturase. A similar mutation in *S. pombe* or *S. cerevisiae* requires exogenous supplementation with oleic acid due to the poor growth of these mutants (*13*–*15*). In contrast, *S. japonicus* cells lacking ole1 grow similarly to the wild-type strain in minimal medium conditions at 37 °C, relying solely on endogenous fatty acid synthesis (Fig. 2A).

**Fig. 2.**
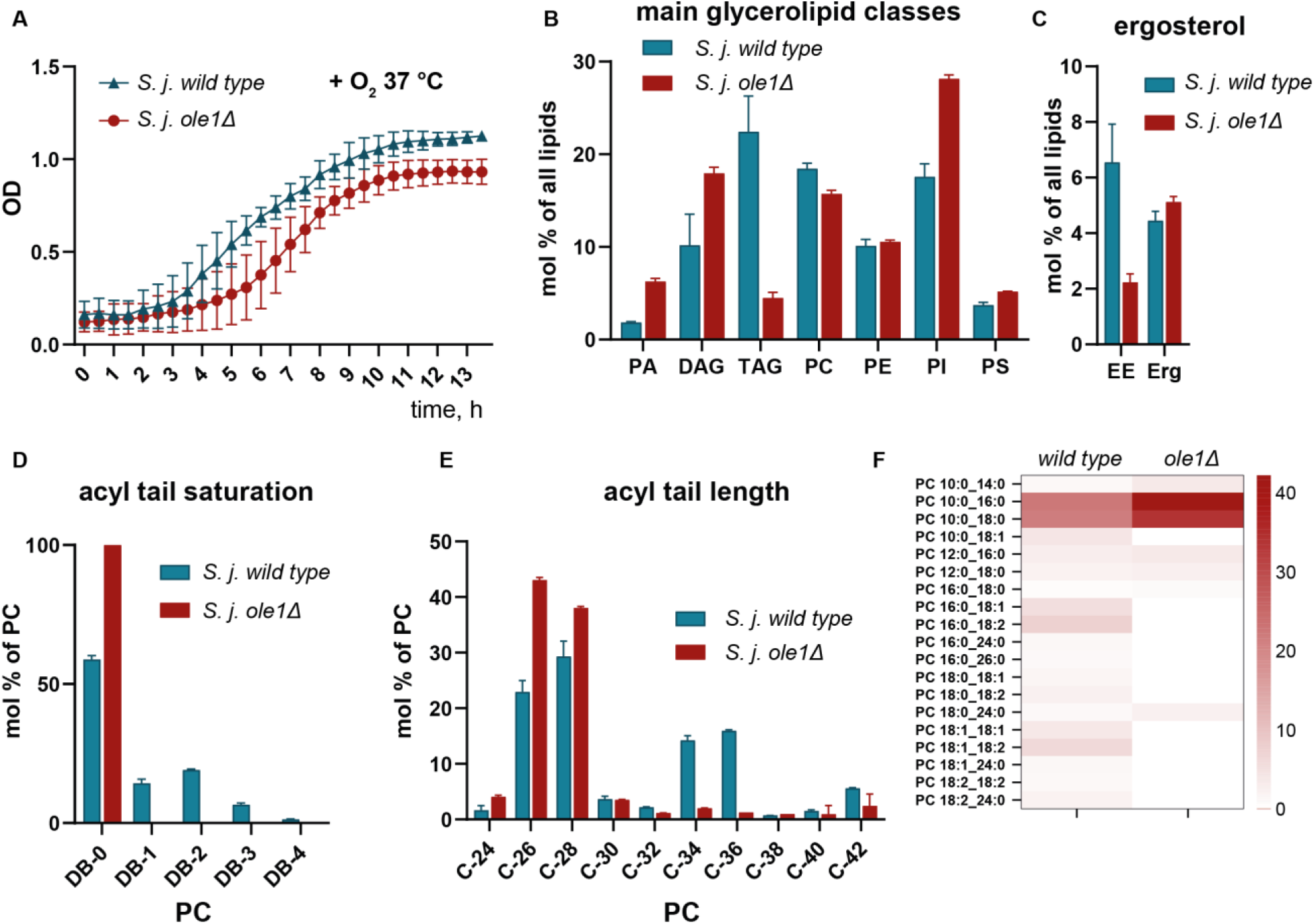
*S. japonicus* lacking desaturase *ole1* also generate saturated, asymmetric phospholipids. (**A**) Growth curves of *S. japonicus* wild type and *S. japonicus ole1Δ* showing OD changes at 37° C. B) Relative abundance of the main lipid classes in *S. japonicus* wild type and *ole1*Δ presented as molecular percentage of identified lipids. C) Relative abundance of ergosterol and ergosterol ester in *S. japonicus* wild type and *ole1*Δ. D) Relative abundance of double bonds in acyl tails within PC class in *S. japonicus* wild type and *ole1*Δ. E) Relative abundance of various carbon lengths in acyl tails in *S. japonicus* wild type and *ole1Δ*. F) A heat map showing molecular species composition within PC class in *S. japonicus* wild type and *ole1*Δ.

To assess alterations to the membrane lipidome in the *ole1Δ* mutant compared to wild type we performed a similar lipidomic analysis where we compared wild type *S. japonicus* cells to *ole1Δ* mutant cells. Quantitative analysis revealed similar changes as were identified in anaerobic conditions. First, as expected, unsaturated lipids were absent in the *ole1Δ* mutant indicating that the deletion was successful in preventing the production of unsaturated phospholipids. Second, the main glycerolipid classes that were significantly increased were PA and DAG, while TAG was decreased (Fig. 2B). Third, and most interesting, the acyl tail length was decreased in a similar manner to that observed for wild type cells grown in anaerobic conditions (Fig. 2D, E and F for PC, the distribution of lipid species in other classes is shown in fig. S2 and table S3, 4).

An important difference between the desaturate deletion mutant and cells grown in anaerobic conditions is that the ergosterol content remained at a similar level as the wild type cells (Fig. 2C). Ergosterol synthesis also requires oxygen (*16*), but it is unaffected by the enzyme deletion. This implies the generation of asymmetric lipids is a compensation mechanism specifically for the absence of unsaturated lipids. It also shows that their presence in the bilayer is not a byproduct of any other metabolic changes that a lack of oxygen might have induced or a consequence of any non-specific effects on cell physiology.

### S. japonicus membrane properties can be maintained at 37 °C even in the absence of Ole1

Cells regulate the lipid composition of membranes to accommodate certain biophysical properties that in turn facilitate membrane-associated cellular properties. Previously it has been shown a) *S. japonicus* membranes are more ordered compared to those of *S. pombe* ones and b) in artificial model membranes, saturated asymmetric lipids can substitute for unsaturated lipids in maintaining membrane lipid order (*17*). We therefore hypothesised that *S. japonicus* membrane order would be maintained even in the absence of the unsaturated lipids usually produced by *ole1Δ* mutant. To investigate this, we used the environmentally sensitive probe di-4-ANEPPDHQ which reports on membrane lipid order through changes in its fluorescence emission – this dye exhibits a red-shifted emission in more disordered membranes. The generalized polarization (GP) parameter, derived from the fluorescence measurements, ranges between +1 and −1, with higher values representing higher lipid order and packing (*18*).

Live-cell imaging indicated that the plasma membranes of *S. japonicus* cells growing aerobically at 37 °C are more ordered than those of *S. pombe* (Fig. 3A, B). Both mechanisms for altering acyl tail composition in *S. japonicus* (genetically, through deletion of *ole1*, and conditionally, through growing cells in an oxygen-free environment) showed smaller changes in lipid bilayer order (Fig. 3C). Deletion of *ole1* resulted in a slight increase in membrane order compared to the wild type. This is consistent with previous results on artificial membranes (*17*). Anaerobically grown cells had a slightly lower membrane order compared to aerobically grown cells, possibly due to a decrease in ergosterol content (Fig. 1E).

**Fig. 3.**
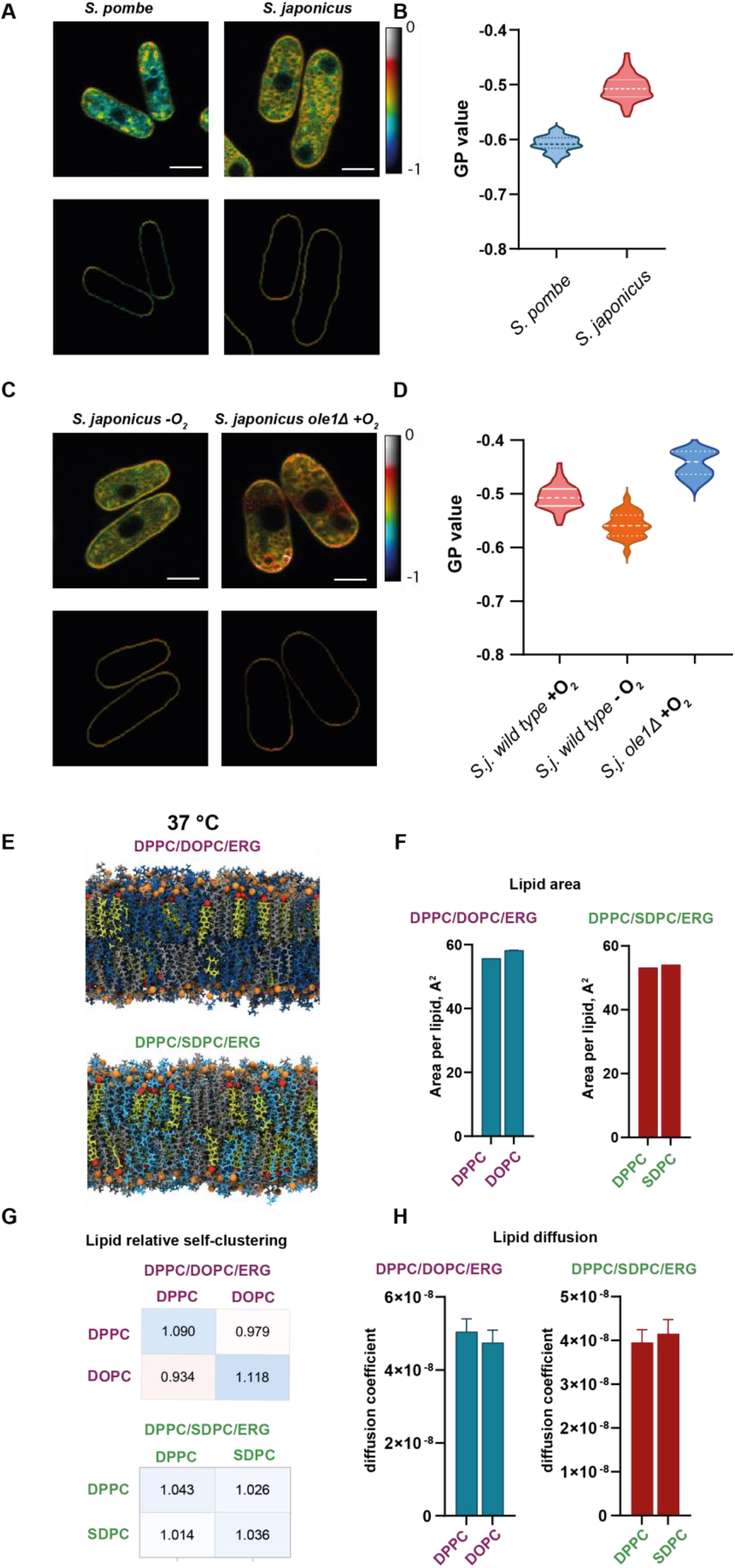
*S. japonicus* lacking desaturase ole1 maintains membrane properties at 37 °C. (**A**) Pseudocoloured generalized polarization (GP) images of *S. pombe* and *S. japonicus* stained with polarity-sensitive dye di-4-ANEPPDHQ (upper panel) and cell outlines were used to quantify GP values of the plasma membrane (lower panel). Scale bar 5μm. (B) GP values quantified from *S. japonicus* (n=53) and *S. pombe* (n=101) grown in aerobic setup cells. (C) Pseudocoloured GP images of *S. japonicus* grown in the absence of oxygen and *ole1Δ* in the presence of oxygen stained with polarity-sensitive dye di-4-ANEPPDHQ (upper panel) and cell outlines were used to quantify GP values of the plasma membrane (lower panel). Scale bar 5μm. (D) Corresponding GP values of *S. japonicus* (n=53) in aerobic condition, *S. japonicus* in anaerobic (n= 28) and *S. japonicus ole1*Δ (n=31). (E) Snapshots of atomistic MD simulations of the two three-component bilayers: symmetric desaturated PC and asymmetric saturated PC. Biophysical properties of bilayers extracted from atomistic MD simulations F-H. (H) Area per lipid, (G) Lipid neighbourhood (H) Lipid diffusion coefficient.

To assess the effect of a lack of unsaturated lipids on other membrane properties, we performed atomistic stale molecular dynamics (MD) simulations. We simulated 3-component lipid bilayers composed of either DPPC (saturated lipid), DOPC (unsaturated lipid) and ergosterol representing the control, wild type condition, or DPPC (saturated lipid), SDPC (saturated, asymmetric lipid) and ergosterol representing the absence of unsaturation, compensated for by asymmetry.In the simulations of the DPPC/DOPC/ergosterol membrane, we find that the DPPC lipids prefer to associate with other DPPC lipids and the DOPC lipids prefer to associate with other DOPC lipids; whereas in the DPPC/SDPC/ergosterol membrane, we find that the DPPC and SDPC lipids are more well mixed (Fig. 3G). However this difference in the distribution of lipids within the membranes does not affect the other physical properties of the lipids we have measured (area per lipid, membrane thickness, lipid diffusion) (Fig. 3F, 3H and fig. S3).

Overall, at 37 °C *S. japonicus* cells grow normally in anaerobic conditions (Fig. 1C) and in the absence of ole1 (Fig. 2A). While both of these conditions result in the loss of unsaturated lipids from the membrane, cells can largely maintain their membrane biophysical properties including membrane lipid order and lipid diffusion. As previously shown in artificial membranes therefore (*17*), saturated, asymmetric phospholipids are able to compensate for the loss of unsaturated lipids caused by either desaturase depletion or oxygen deprivation.

### Asymmetric lipids fail to compensate for the symmetric unsaturated counterparts at temperatures lower than 24 °C

At 37 °C asymmetric lipids, which do not require oxygen for their synthesis, are able to compensate for an absence of unsaturated lipids and closely maintain membrane properties. This raises the question of whether this is a more universal mechanism, and if so, why asymmetry is not well represented across diverse organisms. To answer this question, we investigated whether the *ole1Δ* mutant or anaerobically grown *S. japonicus* can grow at a similar range of temperatures as aerobically grown wild type cells. Although we could not see any significant difference in growth at 37 °C (Fig. 1C, 2A), at 24 °C, the growth of *ole1Δ* mutant cells was significantly suppressed (Fig. 4A). To test whether this was specifically due to the lack of unsaturated lipids and not an unspecific temperature effect, we performed a rescue experiment. Ole1 deficient *S. japonicus* cells grown at 24 °C were supplemented with exogenous oleic acid (18:1). This rescued the growth phenotype, confirming the effect is specifically due to the lack of desaturation (Fig. 4A).

**Fig. 4.**
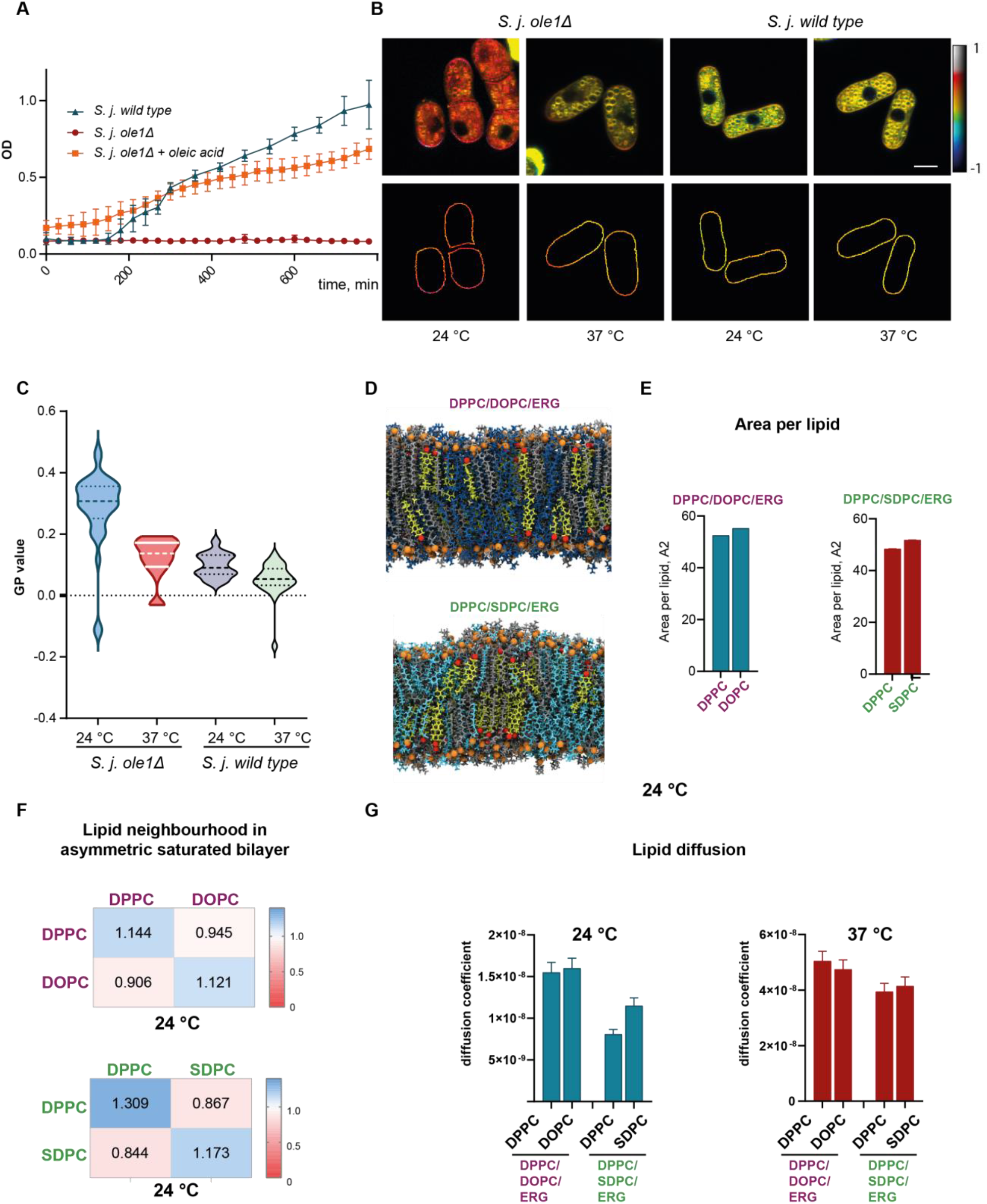
*S. japonicus* lacking desaturase ole1 fails to sustain membrane properties at 24 °C. (**A**) Growth curves of *S. japonicus* wild type and *ole1*Δ, *ole1*Δ supplemented with oleic acid at 24 °C. (B) Pseudocoloured GP images of *S. japonicus* wild type and *ole1*Δ grown at 24 °C and 37 °C and stained with polarity-sensitive dye di-4-ANEPPDHQ (upper panel) and cell outlines used to quantify GP values of the plasma membrane (lower panel). Scale bar 5μm. (C) GP values quantified from n cells. (D) Snapshots of atomistic MD simulations of the two three-component bilayers: symmetric desaturated PC and asymmetric saturated PC. Biophysical properties of bilayers extracted from atomistic MD simulations E-G. (E) Area per lipid, (F) Lipid neighbourhood (G) Lipid diffusion coefficient.

We hypothesised that the failure of ole1 deficient *S. japonicus* cells to grow at 24 °C might be the result of the failure of asymmetric lipids to provide sufficient compensation to maintain membrane properties at this lower temperature. GP values derived from fluorescence microscopy with environmentally-sensitive dyes showed membrane order was increased at 24 °C compared to 37 °C for both wild-type and *ole1Δ* cells (Fig 4B, C), as expected (*17*). However, for wild type cells the effect was extremely small and non-significant. In the *ole1*Δ mutant cells, however, GP values are dramatically increased. This indicated that the cells have failed to maintain membrane fluidity. Molecular dynamics simulations of 3-component lipid bilayers performed at 24°C show little changes in area-per-lipid (Fig. 4E), but self-clustering increases dramatically (Fig. 4D, F). This is particularly true in the system with saturated, asymmetric lipids (DPPC/SDPC/ergosterol) where there is a significant preference for the DPPC lipids to cluster with other DPPC lipids and SDPC lipids to cluster with other SDPC lipids at 24 °C, when at 37 °C, the two lipids were mixed. Most notably, diffusion coefficients only show small differences between 37 °C and 24 °C for bilayers containing unsaturated lipids, but are decreased substantially when unsaturated lipids and replaced by saturated, asymmetric ones (Fig. 4G).

To conclude, while saturated, asymmetric lipids can substitute for unsaturated lipids at 37 °C, this is not the case at 24 °C. At the lower temperature, membrane biophysical properties are compromised, with a loss of fluidity and lipid diffusion. This in turn prevents cell growth, which can be rescued by exogenous supplementation with unsaturated fatty acids (Fig. 4A).

## Discussion

The fluidity of a bilayer is primarily determined by the relative abundance of unsaturated lipids, saturated lipids and sterols, as well as the system’s temperature (*19*). Cells explore and optimise their membrane lipid composition to maintain bilayer physical properties in changing environments (*20*). We have previously shown that novel, saturated, asymmetric phospholipids can substitute for unsaturated lipids in the phase behaviour of ternary artificial membrane systems (*17*). More specifically, a three-component bilayer formed from the asymmetric lipid SDPC together with the symmetric, saturated lipid DPPC and ergosterol will have similar phase and fluidity properties to bilayers composed of symmetric unsaturated lipids (DOPC), DPPC and ergosterol. This may be because the asymmetry generates loser molecular crowding in the hydrophobic core of the bilayer, allowing a greater degree of acyl tail disorder. This leads to the hypothesis that asymmetric phospholipids might allow cells to maintain bilayer fluidity even in the absence of unsaturated lipids.

Here, we demonstrate that this is the case. In response to anoxic conditions, when the desaturase enzymes that produce unsaturated phospholipid tails cannot function, we found that *S. japonicus* produces an abundance of saturated but asymmetric tail phospholipids in response to anoxic conditions. These are capable of replacing symmetric unsaturated lipids in maintaining bilayer biophysical properties. Deletion of the main desaturase *ole1* demonstrated a similar response showing that asymmetric lipids are a more general adaptation to loss of acyl-tail desaturation, rather than lack of oxygen per se. This adaptation allows *S. japonicus* to survive and proliferate in anoxic conditions provided high temperature is maintained. Below 24°C, asymmetric lipids are unable to maintain bilayer fluidity in the same way unsaturated lipids can and growth is curtailed. Conversely, *S. japonicus’* sister species, the well-known model organism *S.pombe*, does not produce an abundance of asymmetric lipids and is incapable of growth in anoxic conditions (Fig. 4C).

The generation of asymmetric lipids to substitute for desaturation in order to maintain membrane fluidity may be a more general adaptation. It is known for example that *S. japonicus* is adapted to natural hypoxic environmental niches. Other organisms that also inhabit these conditions in nature may have developed adaptations to compensate for the difficulty in manufacturing unsaturated membrane lipids. Such information is limited however, primarily because the mass-spectrometry methods to reliably detect asymmetric phospholipids have only recently been widely deployed. Despite this, asymmetric lipids have been detected in *S. cerevisiae* under rich medium conditions (*10*) and there are numerous reports of changes to membrane lipidomes in a variety of hypoxic conditions, for example within the tumour microenvironment or in deep sea marine organisms (*21*–*23*).

By what mechanism can asymmetric phospholipid synthesis be upregulated? A key requirement is the manufacture of medium-chain fatty acids (MCFAs, e.g. C10) to be incorporated into phospholipids. Our previous work has shown that *S. japonicus* is capable of producing MCFAs, most likely through the adaptation of its fatty acid synthase (FAS) enzymes. Purified *S. japonicus* FAS was able to produce MCFAs in vitro whereas *S. pombe* FAS could not (*9*). This might explain the capability of *S. japonicus* in producing MCFAs and hence asymmetric lipids, a capability lacking in *S. pombe*. In order to regulate the length profile of fatty acids produced by FAS, we hypothesise that cells may alter the ratio of acetyl-CoA and malonyl-CoA which are responsible for fatty acid initialisation and lengthening respectively. The full fatty acid length profiles produced by any cell, therefore, is bounded genetically (through FAS) but also modified metabolically e.g. in response to environmental cues.

Despite the insights provided here, several important questions remain unanswered. For example, sterols, of which ergosterol is the most common in fungi also require molecular oxygen for their synthesis (*24*). Since sterols are also important for maintaining membrane fluidity, it is likely that *S. japonicus* may also seek to compensate for the lack of membrane sterols. We hypothesise that hopanoids might represent a candidate substitute for sterols (*25*). It is also unclear what property of the membrane cells sense in order to respond through changes in their lipidome. Our *ole1* deletion experiment shows they are not sensing the lack of oxygen directly, or any other metabolic changes that might be the result of anoxia. However, they may be sensing the absence of acyl-tail desaturation or a property of the bilayer e.g. its lipid packing density or thickness (*26*).

Overall, we have detailed how the ability to synthesise saturated, asymmetric phospholipids allow *S. japonicus* to thrive in anoxic conditions by allowing the organism to maintain the biophysical properties of its membranes. We hypothesise that this might be a more general mechanism for cells and organisms that inhabit hypoxic environmental niches. In general, the adaptation of the membrane lipidome to changing environmental conditions and how lipidomes are genetically and metabolically defined remain poorly understood.

## Acknowledgments

We acknowledge the use of the imaging facility, University of Birmingham.

## Author contributions

Conceptualization: MM

Methodology: LP, CDL, DMO

Investigation: CDL, MM

Visualization: LP, MM

Funding acquisition: MM

Project administration: MM

Supervision: DMO, RCM

Writing – original draft: MM

Writing – review & editing: LP, CDL, RCM, DMO, MM

## Data and materials availability

All strains, custom-reagents and raw data are freely available on request.

