## Supplementary material for "Phospholipid tail asymmetry allows cellular adaptation to anoxic environments": SI

### Materials and Methods

Yeast strains and plasmids

Yeast strains and primers used in this study are listed in Table S1. The deletion of *ole1* gene in *S. japonicus* was performed using a plasmid-based homologous recombination strategy. For that 3'UTR fragment and 5'UTR fragment of *ole1* gene were gene synthesised and cloned in the plasmid with kanR cassette. The plasmid was linearized with SmaI and transformed into wild type *S. japonicus* cells as described previously (*27*). Cells that were successfully transformed were selected on G418 (final concentration 60 µg/ml) for 3 days at 37 °C. The knockout mutant was verified by PCR with the primers listed in Table S1.

Cell growth and media

For genetic modification and cell propagation, *S. japonicus* and *S. pombe* cells were grown in YES medium described in (*28*, *29*). For lipidomic and growth experiments cells were grown Edinburgh minimal medium. For anaerobic experiments, cells were grown initially in aerobic conditions shaking at 170rpm to mid-log growth phase until ~ 0.6OD then shifted to anaerobic cabinet Don Whitley Scientific A35. Cells were diluted to 0.01 OD in a minimal medium that would lack oxygen and grown at 37 °C for 24h. The next day cells were diluted again to 0.01 OD without exposure to atmospheric oxygen and grown for another 24h before any growth, lipidomic and lipid order measurements. This was done to ensure that lipids generated in the presence of oxygen would be used up by the time when samples for lipidomic analysis or growth experiments were collected. For growth experiments cell optical density was measured in aerobic conditions using FLUOstar Omega plate reader at the indicated temperatures in 96 well plates, the measurements were taken every 30 min. For growth measurements in anaerobic conditions, a portable Cerillo Stratus plate reader was used, and cell densities were measured in 6 technical replicates and 3 biological independent replicates in 96 well plates.

Lipid extraction for mass spectrometry lipidomics

Mass spectrometry-based lipid analysis was performed by Lipotype GmbH (Dresden, Germany) as described (*30*, *31*). Lipids were extracted using a two-step chloroform/methanol procedure (*31*). Samples were spiked with internal lipid standard mixture containing: CDP-DAG 17:0/18:1, cardiolipin 14:0/14:0/14:0/14:0 (CL), ceramide 18:1;2/17:0 (Cer), diacylglycerol 17:0/17:0 (DAG), lyso-phosphatidate 17:0 (LPA), lyso-phosphatidylcholine 12:0 (LPC), lyso-phosphatidylethanolamine 17:1 (LPE), lysophosphatidylinositol 17:1 (LPI), lyso-phosphatidylserine 17:1 (LPS), phosphatidate 17:0/14:1 (PA), phosphatidylcholine 17:0/14:1 (PC), phosphatidylethanolamine 17:0/14:1 (PE), phosphatidylglycerol 17:0/14:1 (PG), phosphatidylinositol 17:0/14:1 (PI), phosphatidylserine 17:0/14:1 (PS), ergosterol ester 13:0 (EE), triacylglycerol 17:0/17:0/17:0 (TAG), stigmastatrienol, inositolphosphorylceramide 44:0;2 (IPC), mannosyl-inositolphosphorylceramide 44:0;2 (MIPC) and mannosyl-di-(inositolphosphoryl)ceramide 44:0;2 (M(IP)2C). After extraction, the organic phase was transferred to an infusion plate and dried in a speed vacuum concentrator. 1$^{st}$ step dry extract was re-suspended in 7.5 mM ammonium acetate in chloroform/methanol/propanol (1:2:4, V:V:V) and 2nd step dry extract in 33% ethanol solution of methylamine in chloroform/methanol (0.003:5:1; V:V:V). All liquid handling steps were performed using Hamilton Robotics STARlet robotic platform with the Anti Droplet Control feature for organic solvents pipetting.

MS data acquisition

Samples were analyzed by direct infusion on a QExactive mass spectrometer (Thermo Scientific) equipped with a TriVersa NanoMate ion source (Advion Biosciences). Samples were analyzed in both positive and negative ion modes with a resolution of Rm/z=200=280000 for MS and Rm/z=200=17500 for MSMS experiments, in a single acquisition. MSMS was triggered by an inclusion list encompassing corresponding MS mass ranges scanned in 1 Da increments (*32*). Both MS and MSMS data were combined to monitor EE, DAG and TAG ions as ammonium adducts; PC as an acetate adduct; and CL, PA, PE, PG, PI and PS as deprotonated anions. MS only was used to monitor LPA, LPE, LPI, LPS, IPC, MIPC, M(IP)2C as deprotonated anions; Cer and LPC as acetate adducts and ergosterol as protonated ion of an acetylated derivative (*33*).

MS data analysis and post-processing

Data were analyzed with in-house developed lipid identification software based on LipidXplorer (*34*). Data post-processing and normalization were performed using an in-house developed data management system. Only lipid identifications with a signal-to-noise ratio >5, and a signal intensity 5-fold higher than in corresponding blank samples were considered for further data analysis.

Measurements of membrane lipid order *in vivo*

For imaging, cells were grown in similar conditions for lipidomic analysis and growth experiments, in minimal medium in the presence or absence of oxygen at indicated temperatures overnight and the following day they were diluted to OD 0.2 in the fresh medium containing 5 μM di-4-ANEPPDHQ. Cells were incubated for 2 hours and then transferred into a glass-bottomed microscope dish. Imaging for figure 3 was performed on a Zeiss LSM 780 confocal microscope equipped with a 32 element GaAsP Quasar detector. A 488 nm laser was selected for fluorescence excitation of di-4-ANEPPDHQ. The detection windows were set to 510–580 nm and 620–750 nm. Images from three independent experiments were used for analysis. The setting on the microscope were kept the same between independent experiments. For images in Figure 4 a Zeiss LSM 880 confocal microscope equipped with a 32 element GaAsP Quasar detector with Airyscan was used to obtain images, the settings between sets of experiments were not maintained hence different GP values were obtained between the experiments.

Image analysis

Image pre-processing was undertaken in Fiji image manipulation software version 2.5.0, available under GNU General Public License version 3.0. To increase cell visibility, a copy of each image was created and the auto-contrast plugin was applied. Cell segmentation was then undertaken using the TOBLERONE software package (*35*) written in the R programming language version 4.2.0, employed within the RStudio integrated development environment. Segmentation was achieved by varying the input parameter until the expected number of cells was identified in each image. For each segmented cell, the oriented boundary corresponding to the plasma membrane was then extracted. Generalized polarisation values were calculated on the original images using built-in functions from the TOBLERONE package. Further analysis was undertaken using built-in statistical techniques in R.

Molecular dynamics simulations

Two different lipid bilayers were simulated in order to investigate the effect of lipid molecules with asymmetric tails on the properties of tertiary bilayers where one bilayer consisted of 35% DPPC, 35% DOPC and 30% ergosterol and the other consisted of 35% DPPC, 35% SDPC and 30% ergosterol . Each membrane was built with 200 lipids in each leaflet using the CHARMM-GUI membrane builder (*36*, *37*). Each system also included 40 water molecules per lipid molecule as well as 0.15 mM NaCl. The two bilayers were each studied at two different temperatures, 295 K and 310 K. We performed two replica simulations for each bilayer at each temperature.

Each membrane was minimised and then equilibrated the desired temperature (either 295 K or 310 K) and a pressure of 1 bar following the simulation protocol prescribed by CHARMM-GUI (*38*). After equilibrating each system, a production simulation was carried out for 1 μs at the desired temperature and a pressure of 1 bar. The temperature was controlled by a Nosé-Hoover thermostat and a Parrinello-Rahman barostat was used to control pressure.

All simulations were run using the GROMACS 2020 simulation package (*39*) and the CHARMM36 forcefield was used to model the interactions of the lipid molecules and the ions ((*40*), while the water molecules were modelled using CHARMM TIP3P (*41*). We used the same model for the SDPC lipid, which was generated by truncating the sn-2 tail of the CHARMM36 DSPC model after the 10$^{th}$ carbon, that we used in our previous study of asymmetric lipid membranes (*17*). LINCS constraints were used on the hydrogen-containing bonds in order to allow us to use 2.0 fs timesteps within the production simulations. The last 00 ns of production simulation was used for analysis.

The bilayer properties were characterised by the area per lipid (APL), bilayer thickness, and lipid order parameter ($S_{CD}$) of the lipids within the bilayers. All analysis was conducted using in-house generated python scripts which used MDAnalysis (*42*, *43*) or Lipyphilic (*44*).

A 2D Voronoi tessellation of atomic positions in each leaflet was performed to determine the APL of each component of the bilayer, using the C21 C2 and C31 atoms as seeds for the PC lipids, and O3 atoms for the ergosterol (atoms are given by CHARMM atom names). We calculate the mean APL of each species at each frame, and calculate the standard deviation over time. The lipid order parameter is a measure of the conformational flexibility of acyl chains in a bilayer, and is given by:

$$S_{CD} = \left| \frac{1}{2} \langle 3\, cos^2(\theta) - 1 \rangle \right|$$

where θ is the angle between the bilayer normal and the carbon-hydrogen vector of a carbon atom in an acyl tail, and the average is taken over time and over all molecules of a given species within the membrane. The $S_{CD}$ was calculated for each lipid species as a function of carbon atom position along an acyl chain. Smaller values of $S_{CD}$ indicate a more disordered acyl chain. To calculate the bilayer thickness, we constructed a lateral 6 x 6 grid of the membrane, calculated the mean phosphate-phospahte distance in each grid point, then averaged over the grid. This better accounts for a rough membrane surface, such as with DPPC in the ripple phase. We report the standard devation of the mean membrane thickness over time.

We calculated the lateral mean squared displacement (MSD) of each species using the C2 and O3 atoms of the PC lipids and ergosterol, respectively, over the final 300 ns of each simulation. Before calculating the MSD of each lipid, we first removed the lateral center of mass motion of the bilayer. We then calculated the lateral diffusion coefficient (*D*) for each lipid using the Einstein relation:

$$D = \frac{1}{2d} \lim_{t \to \infty} \frac{\langle r_t - r_{t0} \rangle}{t}$$

where *d* is the system dimension, *r* is the coordinate of the C2 or O3 atom at a given time, *t*, from a time origin, $t_0$. We used the period from 100 ns to 250 ns of the MSD curve to calculate *D* for each individual lipid. We then calculated the average and standard error of *D*.

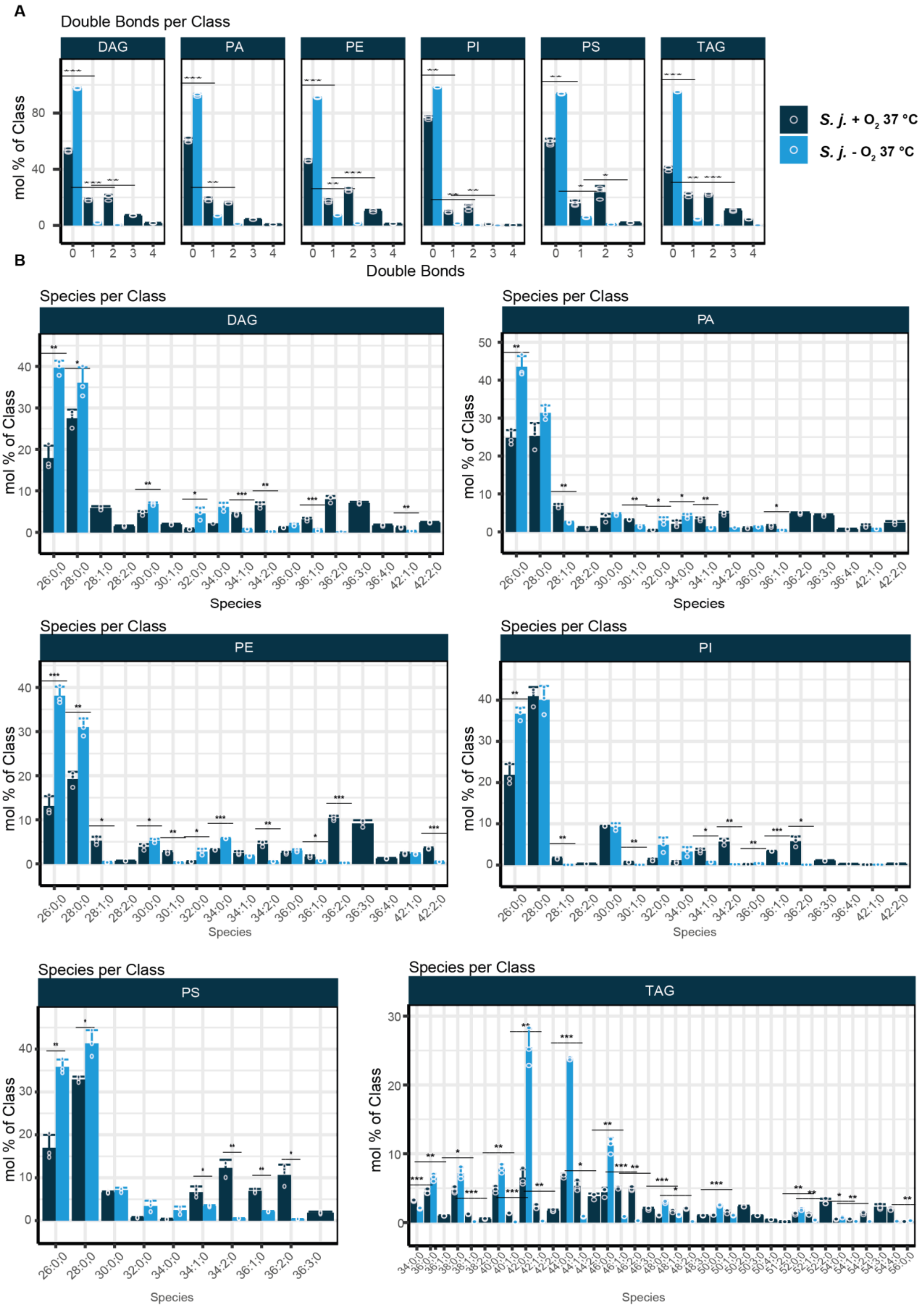
A
Double Bonds per Class
DAG
PA
PE
PI
PS
TAG
mol % of Class
Double Bonds
S. j. + O2 37 °C
S. j. - O2 37 °C
B
Species per Class
Species

**Fig. S1. Quantitative lipidomic results for *S. japonicus* grown in normoxic and anoxic conditions (A)** Relative abundance of double bonds in acyl tails within DAG (diacylglycerol), PA (phosphatidic acid), PE (phosphatidylethanolamine), PI (phosphatidylinositol), PS (phosphatidylserine), and TAG (triacylglycerol) lipid classes in *S. japonicus* grown in normoxic and anoxic conditions. (B) Relative abundance of molecular lipid species within DAG, PA, PE, PI, PS and TAG classes in *S. japonicus* grown in normoxic and anoxic conditions. Presented are mean values of three biological replicates and SD values. T-test was used to indicate significant differences between two conditions.

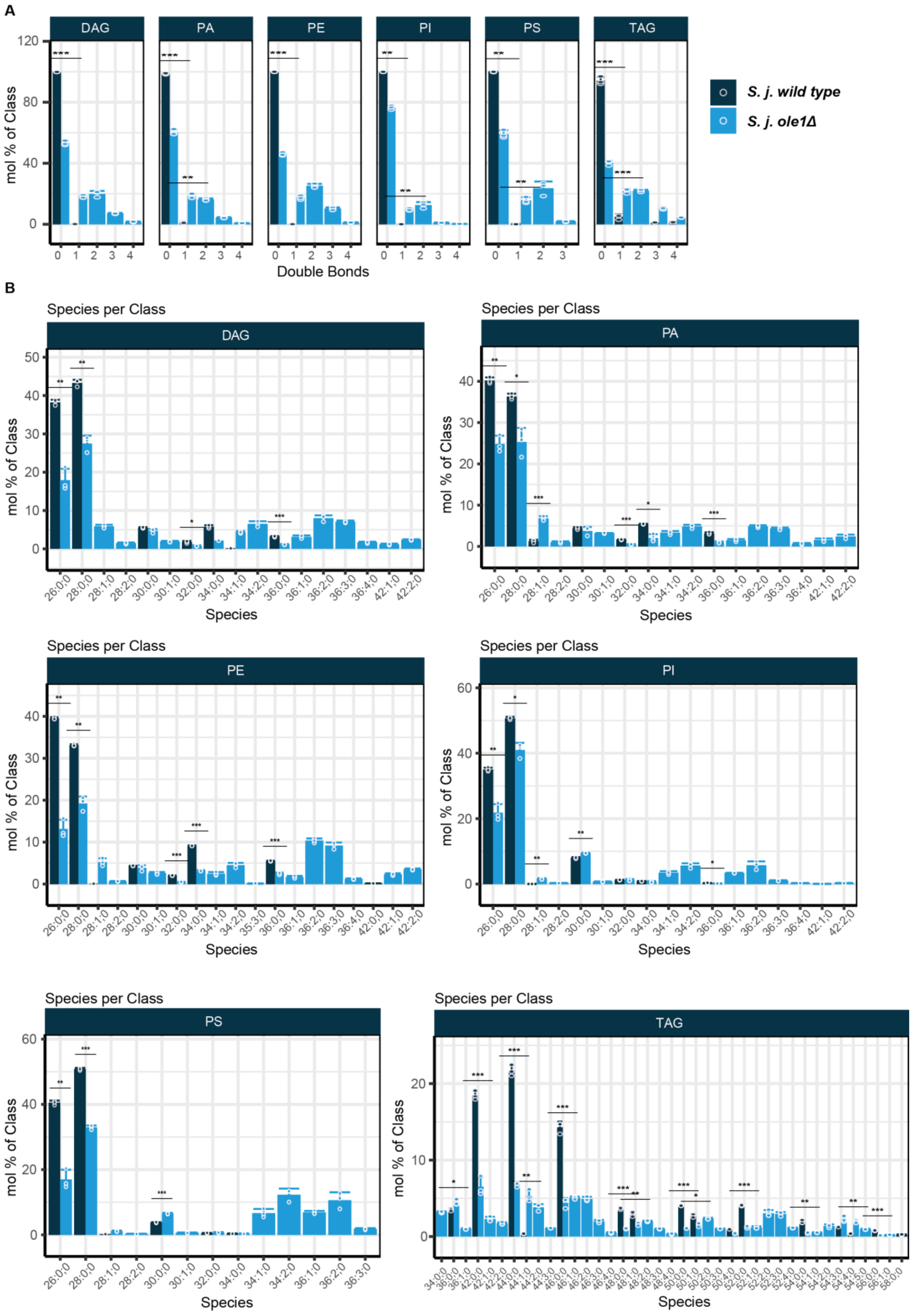
A
DAG
PA
PE
PI
PS
TAG
mol % of Class
Double Bonds
S. j. wild type
S. j. ole1Δ
B
Species per Class
Species

**Fig. S2. Quantitative lipidomic results for *S. japonicus* wild type and *S. japonicus ole1Δ*.** (A) Relative abundance of double bonds in acyl tails within DAG, PA, PE, PI, PS, and TAG lipid classes in *S. japonicus* wild type and *S. japonicus ole1Δ*. (B) Relative abundance of molecular lipid species within DAG, PA, PE, PI, PS and TAG classes in *S. japonicus* wild type and *S. japonicus ole1Δ*. Presented are mean values of three biological replicates and SD values. T-test was used to indicate significant differences between two conditions.

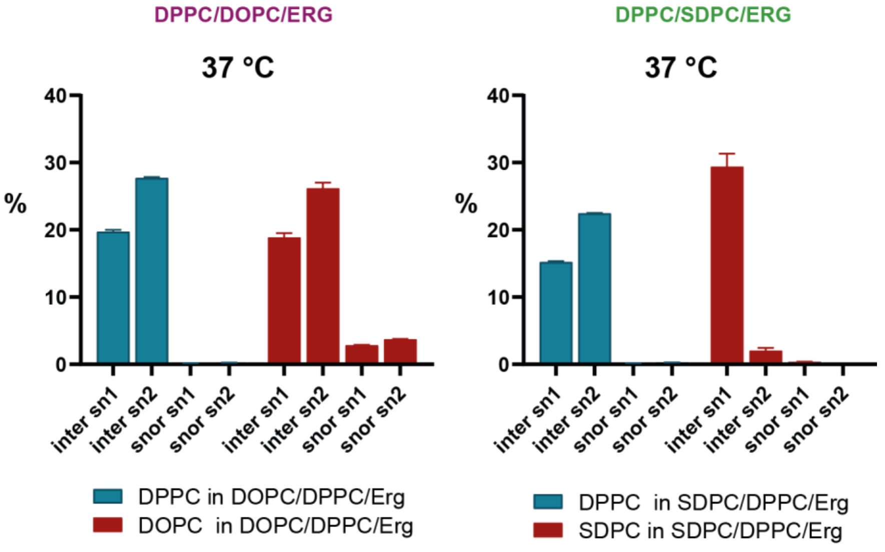


**Fig. S3.** Percentage of the terminal carbons in each of the sn acyl chains that interdigitate or snorkel at 37°C in simulated lipid bilayers composed of either DPPC/DOPC/Erg or DPPC/SDPC/Erg.

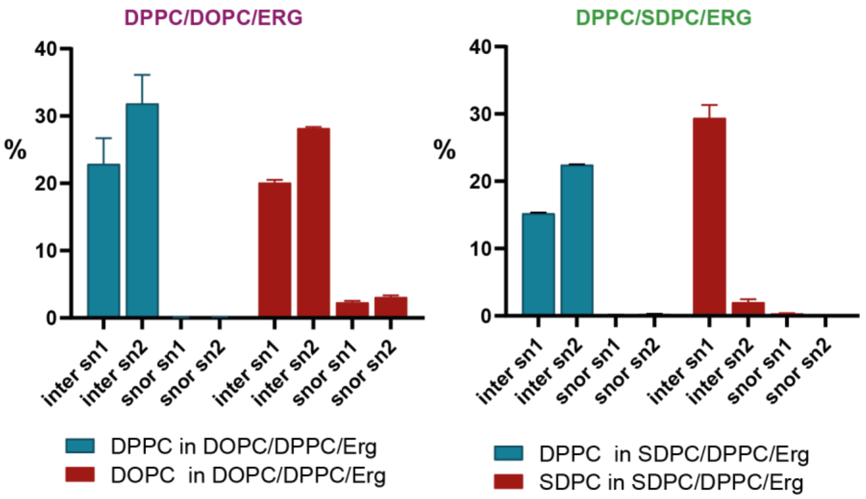


**Fig. S4.** Percentage of the terminal carbons in each of the sn acyl chains that interdigitate or snorkel at 24°C in simulated lipid bilayers composed of either DPPC/DOPC/Erg or DPPC/SDPC/Erg.

**Data S1. (separate file)**

Quantitative lipidomic data
